# Staged heavy-chain filtering enables Fab discovery from combinatorially intractable library spaces

**DOI:** 10.64898/2026.05.10.724059

**Authors:** Youngju Kim, Hayoung Kwon, Jisu Hong, Chang Kyung Kang, Wan Beom Park, Hang-Rae Kim, Chang-Han Lee

## Abstract

**Background:** Combinatorial fragment antigen-binding (Fab) libraries encode an immense heavy–light chain pairing space, often exceeding 10¹⁴ possible combinations, which far surpasses the diversity that can be experimentally constructed and screened in display systems. As a result, direct Fab screening samples only a small fraction of the theoretical search space, creating a practical bottleneck for functional binder discovery.

**Results:** Here, we frame Fab discovery as a staged search problem by decoupling heavy-chain (HC) and light-chain (LC) exploration. We implemented a sequential HC preselection–remating workflow in yeast surface display, in which antigen-reactive HC variants are first enriched and subsequently recombined with a diverse LC repertoire to reconstruct a focused Fab library. In a SARS-CoV-2 spike-targeted campaign, HC and LC libraries of 2.05 × 10⁷ and 2.33 × 10⁷ members corresponded to a theoretical pairing space of approximately 4.8 × 10¹⁴ combinations. Sequential HC enrichment followed by LC remating allowed recovery of multiple functional Fab clones from a tractable library scale of approximately 10⁸, including clones that shared a common HC scaffold but carried distinct LC partners. A representative recombinant IgG output showed broad but heterogeneous spike/RBD binding, measurable pseudovirus neutralization activity (EC₅₀ = 11.1 nM), and compatibility with standard early biophysical characterization after full-length IgG reformatting.

**Conclusions:** These results provide proof of principle that combinatorial Fab discovery can be approached as a staged exploration problem under realistic library-size constraints. By focusing downstream Fab reconstruction on an antigen-compatible HC subspace, sequential HC preselection followed by LC remating offers a practical strategy for exploring otherwise intractable antibody pairing landscapes in eukaryotic display systems.

## Background

Monoclonal antibodies have become a central therapeutic modality across oncology, immunology, and infectious diseases (1–4), driven in part by advances in display-based discovery platforms such as phage, yeast, and mammalian display systems (5–9). These technologies have enabled the interrogation of increasingly large antibody repertoires, substantially expanding the accessible diversity of antigen-binding molecules. Among these approaches, combinatorial fragment antigen-binding (Fab) libraries are particularly attractive because they diversify both heavy-chain (HC) and light-chain (LC) variable regions during primary discovery, thereby broadening the theoretical HC–LC pairing space available for antigen recognition (5).

In principle, independent HC and LC repertoires can encode a theoretical pairing space on the order of 10¹²–10¹⁴ possible combinations (10–13). However, the experimentally accessible diversity of eukaryotic display systems is typically limited to 10⁷–10⁹ variants, representing only a minute fraction of this combinatorial landscape (10,12). Direct Fab screening therefore operates under severe sampling constraints in which the vast majority of potential HC–LC combinations remain unexplored (10,14,15). Compounding this physical limitation, nominally large libraries may still sample HC diversity unevenly, because a limited subset of input HCs can be repeatedly paired with multiple LC partners, leaving potentially valuable HC solutions under-represented before functional selection can act on them.

This challenge is not readily resolved by simply increasing library size or screening throughput alone. Although larger libraries improve sampling density, they do not fundamentally alter the mismatch between the theoretical pairing space and experimentally accessible diversity, because the combinatorial expansion of HC–LC pairings outpaces achievable gains in physical library representation. Efficient strategies for navigating this high-dimensional search space are therefore needed (16).

One potential approach is to reduce the dimensionality of the search space prior to full combinatorial pairing. In many antibody systems, HC complementarity-determining regions (CDRs), particularly CDR-H3, make substantial contributions to antigen recognition and specificity (17,18). This observation raises the possibility that early enrichment of antigen-compatible HC variants could define a functionally relevant subspace for subsequent LC diversification, concentrating downstream Fab reconstruction on HC solutions that have already demonstrated antigen compatibility.

Yeast surface display is well suited to such a staged design. It provides a eukaryotic folding environment compatible with Fab assembly, supports quantitative fluorescence-activated selection, and enables *in vivo* recombination of separately maintained HC and LC repertoires through mating of complementary haploid populations (5,7). These features make it possible to separate HC and LC exploration into sequential stages and then reconstruct a more focused Fab search space after an initial filtering step.

Here, we developed a sequential HC preselection–remating workflow for staged Fab discovery in yeast. HC variants were first screened in a haploid format to enrich antigen-reactive seeds, after which the selected HC pool was remated with a diverse LC repertoire to reconstruct a focused Fab library for downstream selection. Unlike conventional chain-shuffling strategies that are typically applied after a lead antibody has already been identified, this workflow introduces HC-first filtering at the primary discovery stage, before any functional hit is in hand. Thus, the aim is not to refine a known binder but to define a tractable search subspace from which functional Fabs can be initially isolated.

Using SARS-CoV-2 spike protein as a model antigen, we recovered multiple functional Fab clones that shared a preselected HC scaffold but carried distinct LC partners, and validated a representative clone through binding, neutralization, and preliminary biophysical characterization. These results provide proof of principle for sequential HC preselection followed by LC remating as a practical staged strategy for Fab discovery under realistic library-size constraints.

## Materials and Methods

### Isolation and cryopreservation of PBMCs

Peripheral blood mononuclear cells (PBMCs) were isolated from heparinized peripheral whole blood by density-gradient centrifugation using Ficoll-Histopaque (1.077 g/mL; GE Healthcare Life Sciences, Chicago, IL, USA) (19). Purified PBMCs were cryopreserved in liquid nitrogen in freezing medium consisting of 50% fetal bovine serum, 10% dimethyl sulfoxide, and 40% RPMI-1640 (all from Thermo Fisher Scientific, Waltham, MA, USA) until further use.

### Construction of the BCR library from PBMCs of patients with COVID-19

Complementary DNA (cDNA) was synthesized from total RNA extracted from PBMCs of patients with COVID-19. Heavy-chain variable (VH) and light-chain variable (VL) genes were amplified using human antibody-specific primer sets (**Supplementary Table. S1**) and fused to CH1 and CL region genes, respectively, to generate Fab-format gene cassettes (20). The amplified VH-CH1 and VL-CL fragments were separately co-transformed with linearized expression vectors pYDS-H and pYDS-K into *Saccharomyces cerevisiae* strains JAR200 (MATa) and YvH10 (MATα), respectively, by electroporation to enable *in vivo* homologous recombination. The resulting haploid heavy-chain (HC) and light-chain (LC) libraries were inoculated into 300 mL of SDCAA medium supplemented with uracil or tryptophan, respectively, and cultured at 30 °C with shaking at 160 rpm for 15 h until the optical density at 600 nm (OD_600_) reached approximately 5.

Library size was estimated from transformation-derived colony-forming units together with insert-positive rates determined during library quality control. In the SARS-CoV-2 campaign shown in Fig. 1, the HC and LC libraries were estimated to contain 2.05 × 10^7^ ± 0.82 × 10^7^ and 2.33 × 10^7^ ± 0.57 × 10^7^ members, respectively. These values represent the mean ± s.d. of three independent transformations (yielding 1.55 × 10⁷, 1.60 × 10⁷, and 3.00 × 10⁷ CFU for HC, and 2.99 × 10⁷, 2.00 × 10⁷, and 2.00 × 10⁷ CFU for LC). This corresponds to a theoretical HC–LC pairing space of approximately 4.8 × 10¹⁴ combinations. To generate the remated Fab library, an HC pool recovered after prior HC preselection and the LC library were mixed in equal volumes normalized to an OD_600_ of 200. Mating was induced by vortexing and inverting the mixture, followed by incubation on YPD agar plates (pH 4.5) at 30 °C for 8 h. Harvested cells were washed twice with sterile distilled water, resuspended in fresh SDCAA medium to an initial OD_600_ of 0.1, and cultured overnight at 30 °C. Mating efficiency was determined by serial dilution plating onto SDCAA agar and tryptophan-supplemented SDCAA agar, followed by colony counting and calculation of the corresponding ratio.

**Figure 1.**
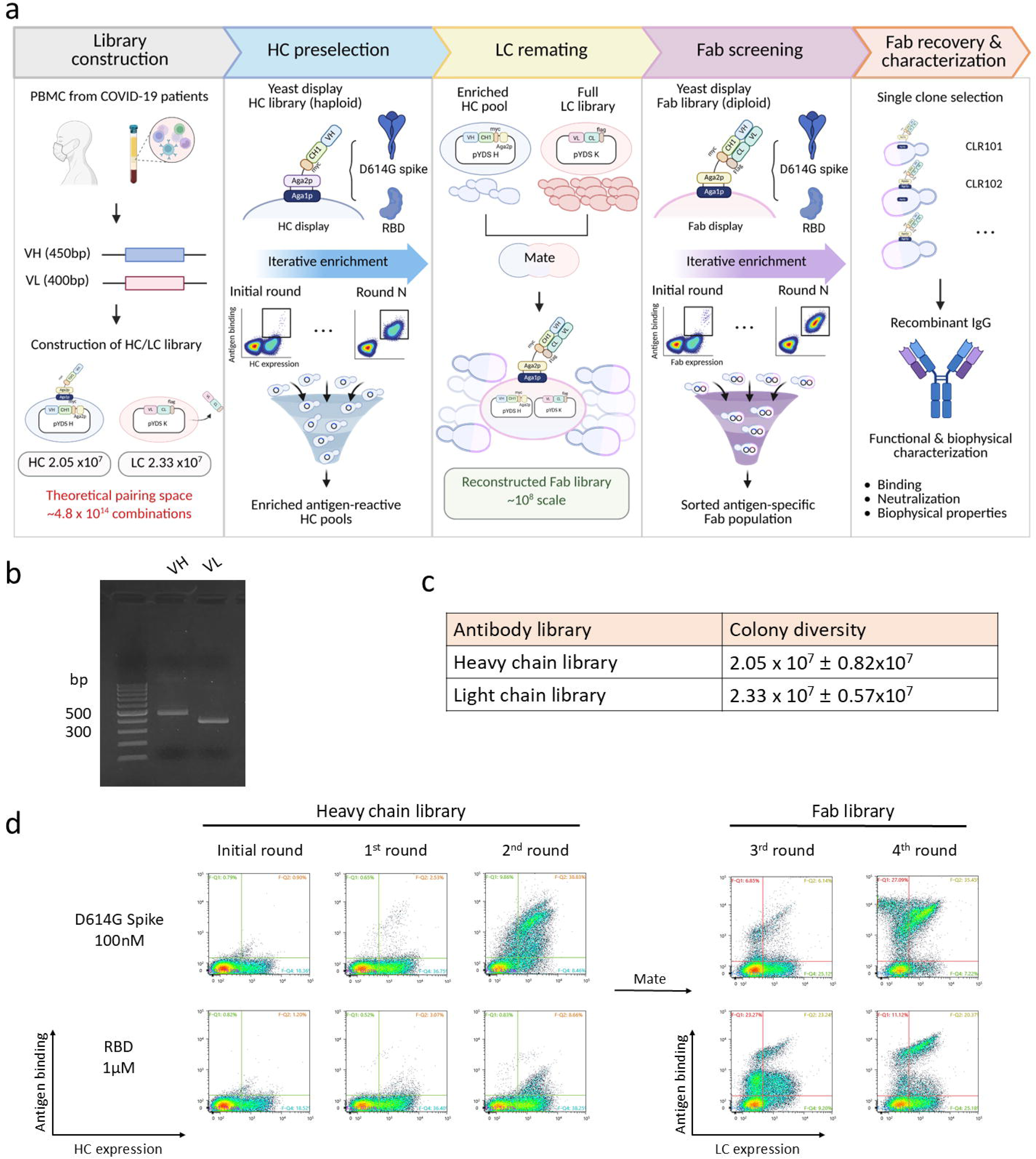
The sequential HC preselection–remating workflow for staged Fab discovery. (a) Schematic representation of the staged Fab discovery platform. (b) PCR amplification of VH (approximately 450 bp) and VL (approximately 400 bp) gene cassettes derived from COVID-19 patients. (c) Estimated colony diversity of the initial HC and LC libraries (n = 3, mean ± s.d.). (d) Flow-cytometric analysis of the antibody screening process. The x-axis represents HC or LC surface expression detected via anti-c-myc antibody (HC library) or anti-FLAG antibody (Fab library), followed by FITC-conjugated anti-mouse IgG secondary antibody, and the y-axis represents antigen binding detected by streptavidin-PE against biotinylated D614G spike protein and biotinylated wild-type RBD. The HC library was first subjected to one round of MACS enrichment followed by one round of FACS sorting against biotinylated D614G spike. Following mating of the selected HC pool with the full LC library (2.33 × 10⁷ members), the reconstructed diploid Fab library was subjected to two rounds of FACS-based sorting against biotinylated wild-type RBD to isolate high-affinity target-specific binders

### Preparation of recombinant SARS-CoV-2 antigens

Genes encoding the SARS-CoV-2 D614G spike protein and the receptor-binding domains (RBDs) of the wild-type (Wuhan-Hu-1), Alpha (B.1.1.7), Beta (B.1.351), Gamma (P.1), Delta (B.1.617.2), and BA.5 (B.1.1.529.5) variants were cloned into the pcDNA3.4 mammalian expression vector using Gibson assembly (New England Biolabs, Ipswich, MA, USA) (19). Recombinant antigens were transiently expressed in Expi293 cells (Thermo Fisher Scientific). Five days after transfection, cell culture supernatants were harvested and His-tagged proteins were purified by affinity chromatography using Ni-NTA agarose resin (QIAGEN, Hilden, Germany), as described previously (19). Supernatants were passed through the Ni-NTA column and washed with 50 mL of wash buffer containing 50 mM sodium phosphate, 300 mM NaCl, and 25 mM imidazole (pH 8.0). Bound proteins were eluted with 15 mL of elution buffer containing 50 mM sodium phosphate, 300 mM NaCl, and 250 mM imidazole (pH 8.0). Buffer exchange into PBS (pH 7.4) was performed using Amicon Ultra-4 centrifugal filter units with a 10-kDa molecular weight cut-off (Merck Millipore, Darmstadt, Germany). Purity and molecular weight of the purified proteins were assessed by 12% SDS-PAGE (**Supplementary Fig. S1a,b**).

### Screening and sorting of yeast surface display libraries

To uniformly monitor the enrichment profile across all selection stages, flow cytometric binding analyses were consistently evaluated using fixed concentrations of 100 nM biotinylated D614G spike protein and 1 µM wild-type RBD. For surface labeling, induced yeast cells were incubated with either anti-c-myc mouse antibody (clone 9E10, 1:100) for HC constructs or anti-FLAG antibody (1:200, Thermo Fisher Scientific; MA1-91878) for Fab constructs, along with the respective biotinylated antigens for 1 h at room temperature. Following washes with PBS containing 0.1% bovine serum albumin (PBSA, pH 7.4), cells were secondarily stained with FITC-conjugated anti-mouse IgG (1:200, Sigma-Aldrich, St. Louis, MO, USA; F0257) and streptavidin-PE (1:100, Thermo Fisher Scientific; S866) for 15 min at 4 °C in the dark. During flow cytometry, single yeast cells were strictly defined by diagonal gating on a forward scatter area (FSC-A) versus forward scatter height (FSC-H) plot to exclude doublets and debris.

Initial enrichment of the HC library was performed by magnetic-activated cell sorting (MACS). Approximately 1 × 10⁹ induced HC-library cells were incubated with biotinylated SARS-CoV-2 D614G spike protein at 1 µM for 1 h at room temperature. After washing with PBSA, cells were resuspended with streptavidin-conjugated magnetic microbeads (Miltenyi Biotec, Bergisch Gladbach, Germany; 130-048-102) according to the manufacturer’s instructions and passed through LS columns on a MidiMACS separator (Miltenyi Biotec). The column was washed 5 times with 10 mL of PBSA, and the retained fraction was eluted by removing the column from the magnetic field and flushing with 6 mL of SDCAA medium. Eluted cells were expanded in SDCAA medium supplemented with uracil at 30 °C overnight. For subsequent flow cytometry-based sorting of the libraries, all procedures were performed using an SH800S cell sorter (Sony Biotechnology Inc., San Jose, CA, USA), with cells gated to strictly isolate the top 1% of antigen-binders (PE) within the expression-positive (FITC) population. For the HC library, cells were labeled exclusively with 100 nM biotinylated D614G spike protein, and a single round of sorting was performed under these conditions. Sorted cells were expanded in SDCAA medium supplemented with uracil for 24 h at 30 °C with shaking at 160 rpm, prior to Fab library reconstruction.

For screening of the remated diploid Fab library, surface expression was induced in 2×SGCAA medium at 20 °C for 48 h. The reconstructed Fab library was subjected to flow cytometry-based sorting using biotinylated wild-type RBD. In the first round, cells were sorted against 100 nM RBD. The sorted cells were then expanded and re-induced for surface expression. Following flow cytometric analysis to confirm enrichment, a second, more stringent sorting round was performed against 1 nM RBD to isolate high-affinity binders. After this final enrichment round, single colonies were isolated from the sorted pool, and colony PCR targeting VH and VL genes was performed to identify the respective HC and LC sequences.

### Expression and purification of selected antibodies

VH and VL fragments from selected clones were subcloned into pCIW-VH and pCIW-VL expression vectors encoding the human IgG1 HC constant region and human kappa LC constant region, respectively, using Gibson assembly. HC and LC plasmids were co-transfected into HEK293F cells at an HC:LC ratio of 1:3. Five days after transfection, culture supernatants were harvested and purified using ProA resin (Amicogen, Jinju, South Korea; Puriose ProA FF-30). After washing with 50 mL of 1× PBS, bound antibodies were eluted with 12 mL of 100 mM glycine (pH 2.5) and immediately neutralized with 1 mL of 1 M Tris-HCl (pH 8.0). Buffer exchange into PBS (pH 7.4) was performed using Amicon Ultra-4 centrifugal filter units with a 30-kDa molecular weight cut-off (Merck Millipore). Purified samples were analyzed by 12% SDS-PAGE (**Supplementary Fig. S1c**).

### Evaluation of antibody binding by ELISA

Antibody binding was evaluated by enzyme-linked immunosorbent assay (ELISA) as previously described (8,21,22). Plates were coated with 50 µL per well of antigen at 2 µg/mL and incubated overnight at 4 °C. After blocking with 3% bovine serum albumin in PBS (PBSB), serially diluted antibodies were added and incubated for 1 h at room temperature. Wells were washed three times with 300 µL of PBS containing 0.05% Tween 20 (PBST). Bound antibodies were detected using HRP-conjugated anti-human IgG Fc antibody (1:12,000, Arigobio, Hsinchu, Taiwan; ARG23874) for 1 h at room temperature. After three additional washes with PBST, the reaction was developed with 3,3′,5,5′-tetramethylbenzidine substrate (TMB; Thermo Fisher Scientific) and stopped with 2 M H_2_SO_4_. Absorbance at 450 nm was measured using an Infinite 200 PRO NanoQuant microplate reader (Tecan Trading AG, Männedorf, Switzerland).

### Evaluation of antibody epitopes by competitive ELISA

Each well of a 96-well MaxiSorp microplate was coated with 50 µL of anti-RBD antibody (2 µg/mL; equivalent to 100 ng/well) and incubated overnight at 4 °C. After blocking with 3% bovine serum albumin in PBS (PBSB) for 1 h at room temperature, 200 nM of biotinylated wild-type (WT) RBD-His antigen alone, or a mixture of biotinylated WT RBD-His (200 nM) and competitor proteins (1 µM of reference anti-RBD antibody or ACE2-mFc) that had been preincubated at 37 °C for 2 h, was added to the wells. The plates were incubated for 1 h at room temperature. Following three washes with 300 µL of PBST, the wells were treated with streptavidin–HRP (1:40,000; Abcam, Cambridge, UK; ab7403) for 1 h at room temperature. After three additional washes with PBST, the reaction was developed with TMB substrate (Thermo Fisher Scientific) and stopped with 2 M H_2_SO_4_. Absorbance at 450 nm was measured using an Infinite 200 PRO NanoQuant microplate reader (Tecan Trading AG). For data analysis, the absorbance signal from the RBD-only control was normalized to 100%, and the competition outcomes were expressed as the percentage of signal reduction relative to the control.

### Pseudovirus production and neutralization assay

Pseudovirus expressing the SARS-CoV-2 S protein was produced as described previously (23,24). SARS-CoV-2 spike-pseudotyped lentiviral particles were generated in HEK293T cells by co-transfection of the HIV-1 NL4-3 ΔEnv Vpr luciferase reporter vector and the GP-pCAGGS plasmid encoding SARS-CoV-2 spike protein. At 48 h after transfection, pseudovirus-containing supernatants were harvested, filtered through a 0.45-µm hydrophilic polyethersulfone membrane (Pall Corporation, Port Washington, NY, USA), and concentrated by ultracentrifugation at 25,700 rpm for 3 h at 4 °C using a Beckman SW32Ti swinging-bucket rotor with thin-wall polypropylene tubes (Beckman Coulter, Brea, CA, USA). Infectious titers were determined by median tissue culture infectious dose (TCID_50_) assay (25).

For neutralization assays, pseudoviruses at 1,300 TCID_50_/mL were pre-incubated with serially diluted monoclonal antibodies for 1 h at 37 °C. Virus–antibody mixtures were then added to monolayers of hACE2-expressing 293T cells (hACE2-293T; Takara Bio Inc., Kusatsu, Shiga, Japan) in 96-well plates. After 48 h of incubation, cells were lysed with 100 µL of 1× luciferase cell culture lysis reagent, and luciferase activity was measured after addition of substrate using a GloMax Discover microplate reader (Promega, Madison, WI, USA). Percent neutralization was normalized by defining uninfected cells as 100% neutralization and cells infected with pseudovirus alone as 0% neutralization. CLR101 was evaluated in parallel with benchmark antibodies CR3022 (26), P2B-2F6 (27), and S309 (28) under the same assay conditions.

### Size-exclusion chromatography analysis

Size-exclusion chromatography (SEC) was performed using a Nexera lite high-performance liquid chromatography system (Shimadzu, Kyoto, Japan) equipped with a Superdex 200 10/300 GL column (10 mm × 300 mm; Cytiva, Marlborough, MA, USA; 28990944). A 20 µg aliquot of antibody at 0.1 mg/mL was injected onto the column pre-equilibrated with PBS (pH 7.4) and eluted at a flow rate of 0.5 mL/min. Elution profiles were monitored by absorbance at 280 nm.

### Intrinsic differential scanning fluorimetry

Thermal unfolding behavior of purified antibodies was assessed by intrinsic differential scanning fluorimetry using a SUPR-DSF (Protein Stable, Leatherhead, Surrey, UK). Samples were prepared in PBS (pH 7.4) at a concentration of 0.5 mg/mL and loaded into 96-well plates. Intrinsic protein fluorescence was monitored at emission wavelengths of 330 nm and 350 nm upon excitation at 280 nm while applying a thermal ramp from 20°C to 105°C at a heating rate of 1°C/min. Apparent unfolding transition temperatures (T_m_) were determined from the maxima of the first-derivative traces of the fluorescence ratio (350 nm/330 nm). All measurements were performed under nonreducing conditions, and each sample was measured in triplicate.

### Statistical analysis and software

All quantitative data are presented as the mean ± standard deviation (s.d.) from at least two or three independent experiments, as specified in the respective figure legends. For enzyme-linked immunosorbent assay (ELISA) and pseudovirus neutralization assays, dose-response curves were generated and the apparent half-maximal effective concentration (EC₅₀) values were calculated using the EC₅₀ Calculator tool provided by AAT Bioquest (AAT Bioquest, Inc., Sunnyvale, CA, USA). Data visualization and graphical plots were generated using GraphPad Prism 9 (GraphPad Software, San Diego, CA, USA). Schematic illustrations, including the graphical abstract and Figure 1a, were created with BioRender.com.

## Results

### Sequential HC preselection and LC remating focus the Fab discovery search space

Conventional direct combinatorial Fab library construction is constrained by the physical limits of yeast transformation and display capacity, which typically restrict experimentally accessible diversity to approximately 10⁷–10⁹ variants (10). In the present study, the SARS-CoV-2 heavy-chain (HC) and light-chain (LC) libraries were estimated to contain 2.05 × 10⁷ ± 0.82 × 10^7^ and 2.33 × 10⁷ ± 0.57 × 10^7^ clones, respectively (**Fig. 1a–c, Supplementary Table S2**). The Cartesian product of these repertoires corresponds to a theoretical pairing space of approximately 4.8 × 10¹⁴ HC–LC combinations, far beyond the scale that can be directly represented and screened in a single yeast-display experiment.

To make this search space experimentally tractable, we implemented a sequential HC preselection–remating workflow rather than attempting to reconstruct the full HC–LC combinatorial landscape at the outset. The HC-only display population was first subjected to staged enrichment to identify antigen-compatible HC seeds prior to LC pairing. One round of magnetic-activated cell sorting followed by two rounds of fluorescence-activated cell sorting progressively enriched spike-reactive HC populations (**Fig. 1d**). The enriched HC pool was then remated with the full LC library. This remating step proceeded with a measured mating efficiency of 66.32%, yielding a reconstructed diploid Fab population of approximately 1.41 × 10⁸ ± 2.30 × 10⁷ cells, a scale compatible with downstream yeast-display sorting.

Subsequent Fab-level sorting of this reconstructed diploid population recovered a distinct antigen-positive population (**Fig. 1d**). These results show that staged HC enrichment followed by LC remating can concentrate downstream Fab reconstruction on an antigen-enriched HC subspace, generating a focused Fab library within practical yeast-display limits rather than attempting to directly sample the full 4.8 × 10^14^ combinatorial landscape.

Together, these findings support the feasibility of sequential HC preselection followed by LC remating as a staged search strategy for Fab discovery under realistic library-size constraints. However, because a matched direct Fab library constructed from the same starting repertoires was not screened in parallel, the present data should be interpreted as a proof-of-principle demonstration of tractability rather than as a direct comparison with conventional Fab screening.

### Light-chain remating recovers multiple Fab outputs from a shared preselected heavy chain

Single clones were isolated from the highest antigen-positive fraction within the expression-positive gate after the final round of reconstructed Fab screening and were subjected to colony PCR and sequence analysis. Among the sequenced clones, three representative examples—CLR101, CLR102, and CLR103—shared a common HC scaffold, including an identical CDR-H3 sequence, but carried distinct LC sequences (**Fig. 2a and Supplementary Table S3**). This result indicates that LC remating recovered multiple paired Fab outputs from a shared preselected HC background.

**Figure 2.**
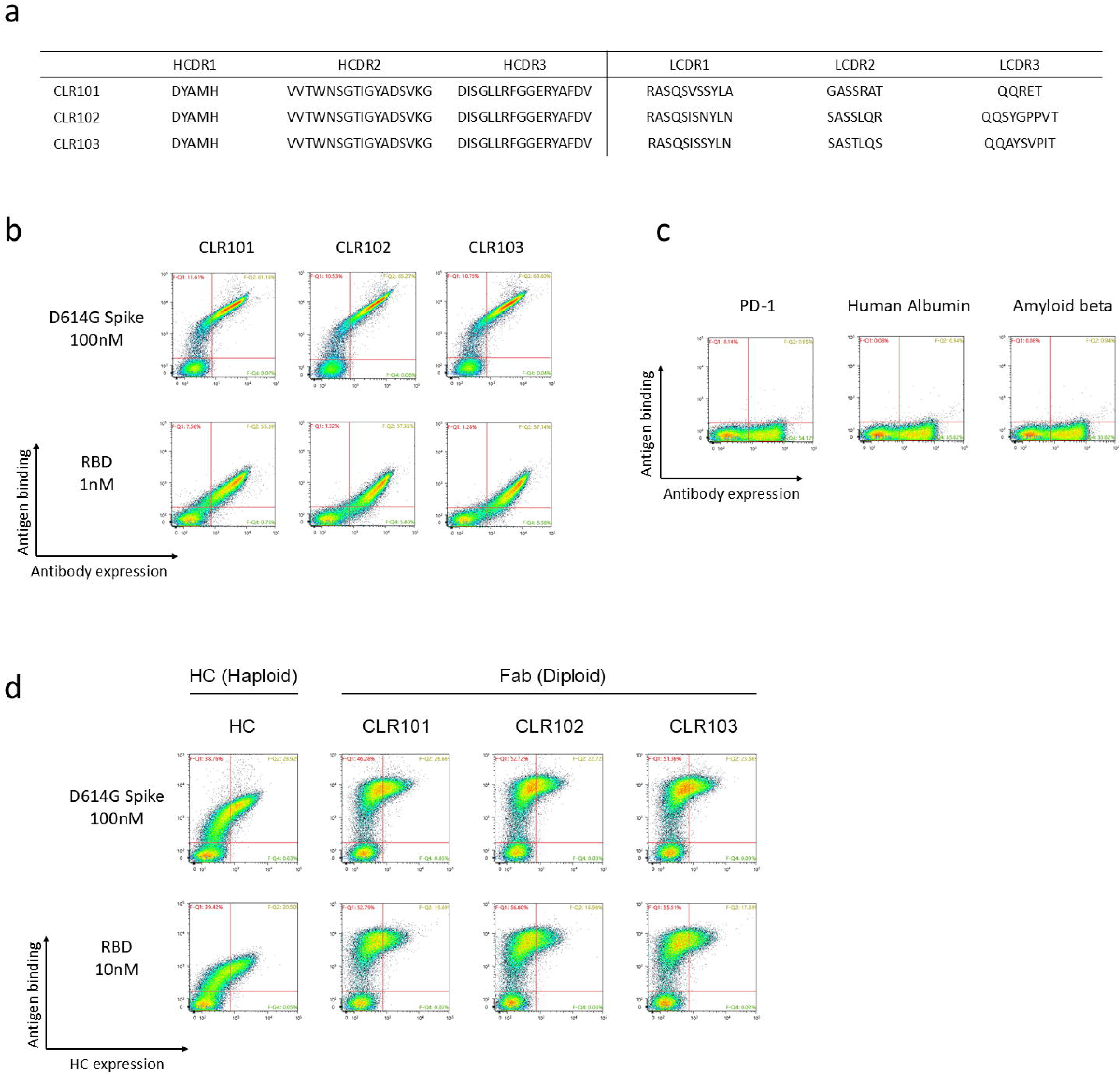
Characterization of remated Fab clones sharing a common heavy-chain scaffold. (a) Amino acid sequences of the complementarity-determining regions (CDRs) for three representative remated Fab clones recovered from the final sorting-enriched population: CLR101, CLR102, and CLR103. (b) Flow cytometry binding analysis of the fully assembled Fab clones against 100 nM D614G spike and 1 nM RBD antigens. (c) Specificity profiling of CLR101 Fab against unrelated antigens (each at 100 nM), including biotinylated PD-1, human albumin, and amyloid beta. (d) Comparison of target binding between the single-chain HC (haploid) format and fully assembled Fab (diploid) formats. Both cell formats were evaluated for binding against 100 nM D614G spike protein and 10 nM wild-type RBD. The Fab format showed higher raw binding signals than the HC-only format; average MFI values increased from 6,037 in the HC-only format to 18,100, 20,800, and 19,300 for CLR101, CLR102, and CLR103 against D614G spike, and from 1,921 to 14,700, 12,900, and 13,500 against wild-type RBD, respectively. Because surface expression levels and display density may differ between formats, these differences should not be interpreted as direct evidence of affinity enhancement.

Flow-cytometric binding analysis showed that all three remated Fab clones bound both D614G spike and wild-type RBD in the assembled Fab format (**Fig. 2b**). To assess whether CLR101 exhibited non-specific binding, the representative clone was tested against unrelated antigens, each at 100 nM, including PD-1, human albumin, and amyloid beta. CLR101 showed no detectable binding to any of the unrelated antigens tested (**Fig. 2c**), supporting target-selective binding within the specificity panel examined here.

We next examined whether target binding was already detectable at the HC stage prior to LC pairing. The HC-only haploid format showed detectable binding to both D614G spike and wild- type RBD, consistent with the recovery of an antigen-compatible HC variant during preselection (**Fig. 2d**). All three Fab clones showed higher antigen-binding signals than the HC-only format (**Fig. 2d**). However, because the HC-only and Fab formats may differ in surface expression level, folding efficiency, or display density, the increase in raw mean fluorescence intensity should not be interpreted as direct evidence of affinity enhancement. This difference should therefore be interpreted with caution in the absence of expression-normalized controls.

Together, these data support the feasibility of using HC preselection to recover antigen-compatible HC variants and subsequent LC remating to generate multiple functional paired Fab outputs from that filtered subspace. The convergence of the analyzed clones on a shared HC background suggests successful enrichment of at least one dominant antigen-compatible HC lineage, while the recovery of distinct LC partners indicates that downstream LC diversification remained possible after HC filtering. Because the present clone-level analysis does not quantify the full diversity of the preselected HC pool or the remated Fab output, the extent to which this workflow preserves broader HC diversity will require deeper sequencing-based analysis.

### CLR101 is a representative recombinant IgG output with broad but heterogeneous SARS-CoV-2 binding

We selected CLR101 as a representative clone for orthogonal characterization after reformatting into full-length human IgG1. ELISA analysis against full-length D614G spike and a panel of variant RBD antigens showed that CLR101 retained binding across the tested SARS-CoV-2 antigens, including WT, Alpha, Beta, Gamma, Delta, and BA.5-derived RBDs (**Fig. 3a–c**).

**Figure 3.**
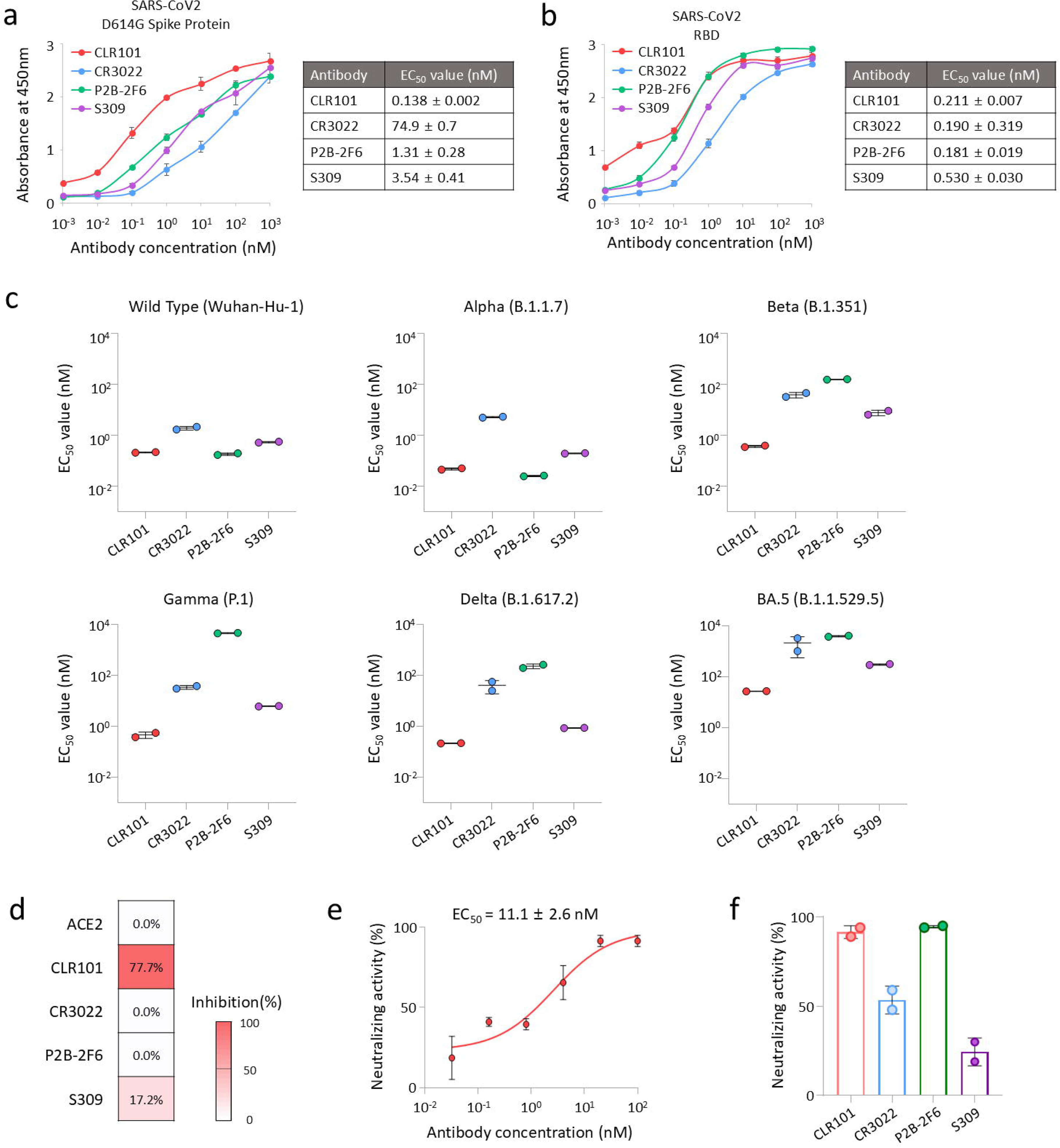
Binding breadth and neutralizing activity of CLR101 selected via the sequential screening workflow. (a, b) Binding ELISA curves and apparent half-maximal effective concentration (EC₅₀) values of CLR101 and reference antibodies (CR3022, P2B-2F6, and S309) against the SARS-CoV-2 D614G spike protein (a) and wild-type RBD (b) (n = 2 independent experiments, mean ± s.d.). (c) Apparent EC₅₀ values of CLR101, CR3022, P2B-2F6, and S309 against RBDs of six SARS-CoV-2 variants—wild-type (Wuhan-Hu-1), Alpha (B.1.1.7), Beta (B.1.351), Gamma (P.1), Delta (B.1.617.2), and BA.5 (B.1.1.529.5)—determined by ELISA (n = 2 independent experiments, mean ± s.d.). (d) Epitope binning analysis of CLR101 by competitive ELISA. The heatmap displays the mean percent inhibition of CLR101 binding to wild-type RBD in the presence of excess competitor proteins (ACE2, CLR101, CR3022, P2B-2F6, and S309). Self-competition by CLR101 was included as a positive control for binding inhibition. The color scale indicates the degree of inhibition from 0% to 100% (n = 3 independent experiments). (e, f) Evaluation of *in vitro* neutralizing activity against D614G spike-pseudotyped lentiviral particles using hACE2-293T cells. (e) Dose-response neutralization curve of CLR101, showing neutralizing activity with an apparent EC₅₀ of 11.1 ± 2.6 nM (n = 2 independent experiments, mean ± s.d.). (f) Percent neutralizing activity of CLR101 alongside benchmark antibodies at a fixed antibody concentration of 100 nM (n = 2 independent experiments, mean ± s.d.).

Against D614G spike, CLR101 showed an apparent EC₅₀ of 0.138 ± 0.002 nM, compared with 74.9 ± 0.7 nM for CR3022, 1.31 ± 0.28 nM for P2B-2F6, and 3.54 ± 0.41 nM for S309 (Fig. 3a). The comparatively high apparent EC₅₀ of CR3022 against full-length spike is consistent with the limited accessibility of its cryptic RBD epitope in the prefusion spike conformation (29). Against wild-type RBD (Wuhan-Hu-1), CLR101 showed an apparent EC₅₀ of 0.211 ± 0.007 nM, comparable to P2B-2F6 (0.181 ± 0.019 nM), whereas S309 showed an apparent EC₅₀ of 0.530 ± 0.030 nM under the same assay conditions (**Fig. 3b**). CR3022 also bound wild-type RBD with an apparent EC₅₀ of 0.190 ± 0.319 nM, although this estimate should be interpreted with caution because of the limited replicate number and high replicate-to-replicate variability. Across the variant RBD panel, CLR101 maintained detectable binding to all tested variants, although the magnitude of binding varied. Binding to WT, Alpha, Beta, Gamma, and Delta remained in the sub-nanomolar to low-nanomolar apparent EC₅₀ range, whereas binding to BA.5 was weaker, with an apparent EC₅₀ of 26.9 ± 0.5 nM (**Fig. 3c, Supplementary Fig. S2a–e, and Supplementary Table S4**) (30). These results indicate that CLR101 has broad but heterogeneous RBD reactivity, with reduced binding to the BA.5-derived antigen relative to earlier variants.

To assess the antigenic topology of CLR101 binding, we performed competitive ELISA using ACE2 and reference RBD-targeting antibodies CR3022, P2B-2F6, and S309 (**Fig. 3d**). ACE2 did not inhibit CLR101 binding to RBD, indicating that CLR101 does not primarily target the receptor-binding motif. CLR101 also showed little to no competition with CR3022 or P2B-2F6. Partial competition was observed with S309, resulting in 17.2% inhibition, although this modest reduction should be interpreted cautiously given the limited replicate number and assay variability. Unlabeled CLR101 produced 77.7% inhibition as a self-competition control. These results suggest that CLR101 recognizes an epitope topologically distinct from the ACE2-binding site and from the binding sites of CR3022 and P2B-2F6, with possible partial overlap or steric proximity to the S309-recognized epitope.

We next evaluated whether CLR101 retained functional activity in a D614G spike-pseudotyped neutralization assay. CLR101 showed neutralizing activity with an apparent EC_50_of 11.1 ± 2.6 nM (**Fig. 3e**). At a fixed antibody concentration, CLR101 and P2B-2F6 achieved 91.5% and 94.5% neutralization, respectively, whereas CR3022 and S309 reached approximately 53.5% and 25% neutralization under the same assay conditions (**Fig. 3f**). Because the activity of benchmark antibodies can vary depending on pseudovirus format, spike density, antibody preparation, and assay conditions, these comparator data should be interpreted within the context of this assay rather than as a general potency ranking. Together, these findings indicate that CLR101 is a functional recombinant IgG output of the sequential HC preselection–remating workflow, with broad but heterogeneous SARS-CoV-2 binding and measurable neutralizing activity.

### Early biophysical characterization supports tractable behavior of the representative clone

We next examined the biophysical properties of CLR101 following reformatting into full-length human IgG1. SDS-PAGE confirmed appropriate expression and assembly of the recombinant antibody (**Fig. 4a**). Size-exclusion chromatography showed a dominant monomer peak with no detectable high-molecular-weight species under the tested conditions (**Fig. 4b**). Intrinsic differential scanning fluorimetry revealed two unfolding transitions with apparent Tm values of 73.5 °C and 77.9 °C (**Fig. 4c and Supplementary Table S5**).

**Figure 4.**
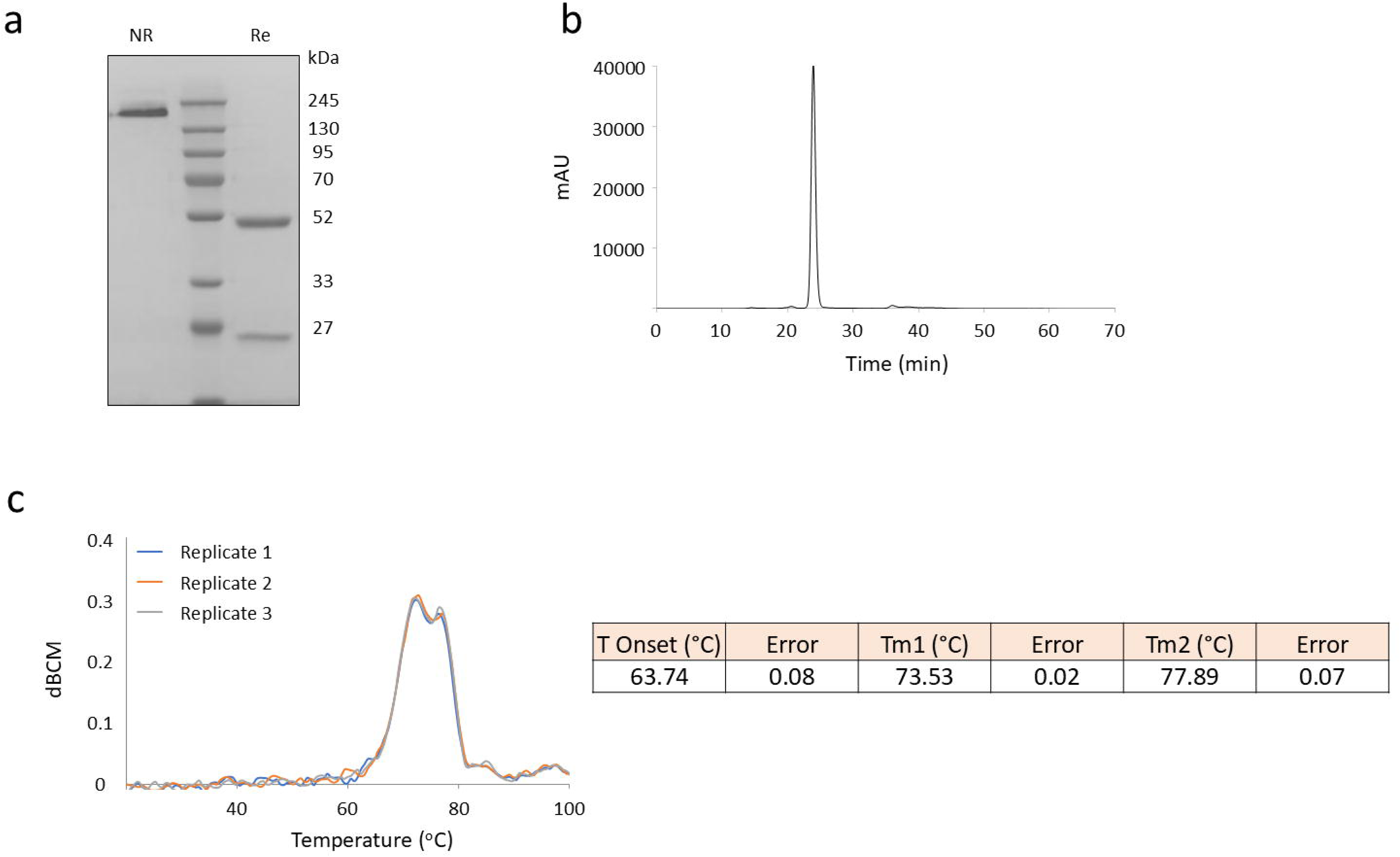
Biophysical characterization of CLR101 IgG1. (a) SDS-PAGE analysis of purified CLR101 under non-reducing (NR) and reducing (Re) conditions. (b) Size-exclusion chromatography (SEC) profile showing a dominant monomeric peak. (c) Intrinsic differential scanning fluorimetry, also referred to as SUPR-DSF, measuring the thermal unfolding transitions (Tm) of the antibody (n = 3 independent replicates). Three replicates show reproducible first-derivative traces of the fluorescence ratio (350 nm/330 nm).

Together, these results indicate that the representative clone recovered through the sequential HC preselection–remating workflow can be reformatted into a full-length IgG and subjected to standard early biophysical characterization.

## Discussion

A central challenge in combinatorial Fab discovery is that productive antigen recognition emerges from paired heavy-chain (HC) and light-chain (LC) variable regions, whereas experimentally accessible library size is constrained by transformation efficiency and screening throughput (5,10–12). In direct Fab screening, this mismatch creates a substantial sampling bottleneck because only a small fraction of the theoretical HC–LC pairing landscape can be physically represented and interrogated in practice (10–12,14). In the present study, we addressed this challenge by restructuring Fab discovery as a staged search process. Rather than attempting to screen the full combinatorial Fab landscape at once, we first enriched antigen-compatible HC seeds and then reconstructed a focused Fab library by remating those HC seeds with a diverse LC repertoire. This design principle, concentrating functional selection on a pre-filtered subspace rather than the full combinatorial landscape, may be generalizable beyond the specific antigen and library system used here.

A key finding of this study was the recovery of multiple functional Fab variants, CLR101, CLR102, and CLR103, that shared a common preselected HC background but carried distinct LC partners. This result supports the feasibility of HC preselection followed by LC remating as a staged strategy for generating paired Fab outputs from a filtered search space. The convergence of the analyzed clones on an identical CDR-H3 sequence is consistent with the dominant contribution of CDR-H3 to antigen recognition in many antibody responses (17,18,31). At the same time, the recovery of distinct LC partners indicates that downstream LC diversification remained possible after HC filtering. Thus, the workflow did not simply isolate a single HC-only binder, but allowed reconstruction of multiple paired Fab solutions from a shared HC seed.

The role of LC remating may extend beyond simple restoration of Fab assembly. Because the recovered clones shared a common HC background yet carried distinct LCs, alternative LC partners may influence apparent binding strength, paratope presentation, epitope fine-specificity, or antigen-contact geometry (32). In this view, LC remating functions as a second-stage diversification step that can generate related but nonidentical recognition solutions from a conserved HC seed. This interpretation is consistent with prior studies showing that LC pairing can substantially alter antibody binding behavior when paired with a fixed HC (5,33–35).

The present workflow is conceptually distinct from conventional post-hit chain-shuffling strategies. Chain shuffling has commonly been used after a lead antibody has already been identified, primarily for affinity maturation, specificity tuning, or focused optimization (5,35). By contrast, the sequential HC preselection–remating strategy introduces HC-first filtering at the primary discovery stage, before a final Fab hit is available. The goal is therefore not to optimize an existing antibody, but to define a tractable search subspace from which functional Fab binders can be initially recovered. This distinction positions the workflow between direct combinatorial Fab screening and post-hit chain-shuffling: it retains the ability to generate paired Fab outputs while reducing the search burden before full HC–LC recombination.

Yeast surface display is particularly well suited to this staged design. It provides a eukaryotic folding environment compatible with Fab assembly, supports quantitative fluorescence-based enrichment, and enables mating between complementary haploid strains to recombine separately maintained HC and LC repertoires *in vivo* (5,7,10). These features allow HC enrichment and Fab reconstruction to occur within a single display ecosystem, avoiding the need to transfer selected repertoires between unrelated platforms. Accordingly, the contribution of this study is both conceptual and methodological: it demonstrates that yeast mating can serve as an experimentally tractable bridge between early chain-level filtering and downstream Fab-level discovery.

Several limitations should be considered. First, this study did not include a matched direct Fab library generated from the same starting repertoires. Therefore, the present data demonstrate feasibility and tractability of the staged workflow, but do not provide a direct quantitative comparison with conventional Fab screening. Second, the clone-level analysis sampled only a limited fraction of the preselected HC pool and remated Fab output. Deeper sequencing of both populations will be needed to determine how broadly HC diversity is preserved after preselection and whether the observed convergence reflects a restrictive bottleneck or productive enrichment of a dominant antigen-compatible lineage, as convergent antibody responses can emerge during antigen-driven selection (36). Third, the comparison between HC-only and Fab formats was based on raw flow-cytometric binding signals. Because surface expression, folding efficiency, and display density may differ between these formats, the increased binding signal observed after LC pairing should not be interpreted as direct evidence of affinity enhancement. Expression-normalized binding analysis and irrelevant LC controls will be required to separate LC-mediated binding effects from display-related differences. Fourth, binding characterization relied primarily on ELISA-derived apparent EC₅₀ values, and orthogonal kinetic measurements such as bio-layer interferometry or surface plasmon resonance will be needed to define association and dissociation kinetics. Similarly, the modest competitive inhibition observed with S309 (17.2%) was based on a limited number of replicates, and its distinction from assay background variability remains to be confirmed. Although competitive ELISA provided an initial epitope-topology profile for CLR101, comparative cross-competition or structural analyses among sibling clones will be required to determine whether LC variation produces measurable epitope shifts. Finally, CLR101 was used as a representative recombinant IgG output to validate downstream functionality, but further structural and *in vivo* studies would be required to assess its mechanism and translational potential as an antibody candidate.

Taken together, our findings provide proof of principle that combinatorial Fab discovery can be approached as a staged search problem under realistic library-size constraints. Sequential HC preselection followed by LC remating allows downstream Fab reconstruction to focus on an antigen-enriched HC subspace rather than attempting to directly sample the full theoretical HC–LC pairing landscape. This strategy may be particularly useful in eukaryotic display settings where full combinatorial Fab construction is limited by physical library-size constraints and where early concentration on antigen-compatible HC seeds can make functional Fab discovery more experimentally tractable.

## Conclusions

Despite these limitations, our results provide proof of principle that combinatorial Fab discovery can be framed as a staged exploration problem under realistic library-size constraints. By using HC preselection to define an antigen-compatible subspace for subsequent LC diversification, the sequential HC preselection–remating workflow offers a practical strategy for functional antibody discovery when exhaustive exploration of the full HC–LC pairing landscape is experimentally infeasible.

More broadly, this approach may be particularly useful in eukaryotic display systems in which full combinatorial Fab construction is limited by physical library-size constraints, and in discovery settings where early concentration on antigen-compatible HC variants facilitates more tractable downstream Fab reconstruction.

## Ethics approval and consent to participate

This study was approved by the Institutional Review Board of Seoul National University Hospital (IRB No. H-2005-004-1121). Written informed consent was obtained from all participants prior to blood collection.

## Consent for publication

Not applicable.

## Acknowledgements

Not applicable.

## Authors’ contributions

Y.K. designed and performed experiments, analyzed data, and wrote the manuscript. H.K. and J.H. contributed to the construction of antibody and antigen expression vectors and protein purification. C.K.K. and W.B.P. contributed to patient sample collection, clinical coordination, and interpretation of SARS-CoV-2-related immunological materials. H.-R.K. contributed to study design, clinical sample resources, and manuscript review. C.-H.L. conceived and supervised the study, secured funding, interpreted data, and revised the manuscript.

## Availability of data and materials

All data that support the findings of this study are available from the corresponding author upon reasonable request.

---

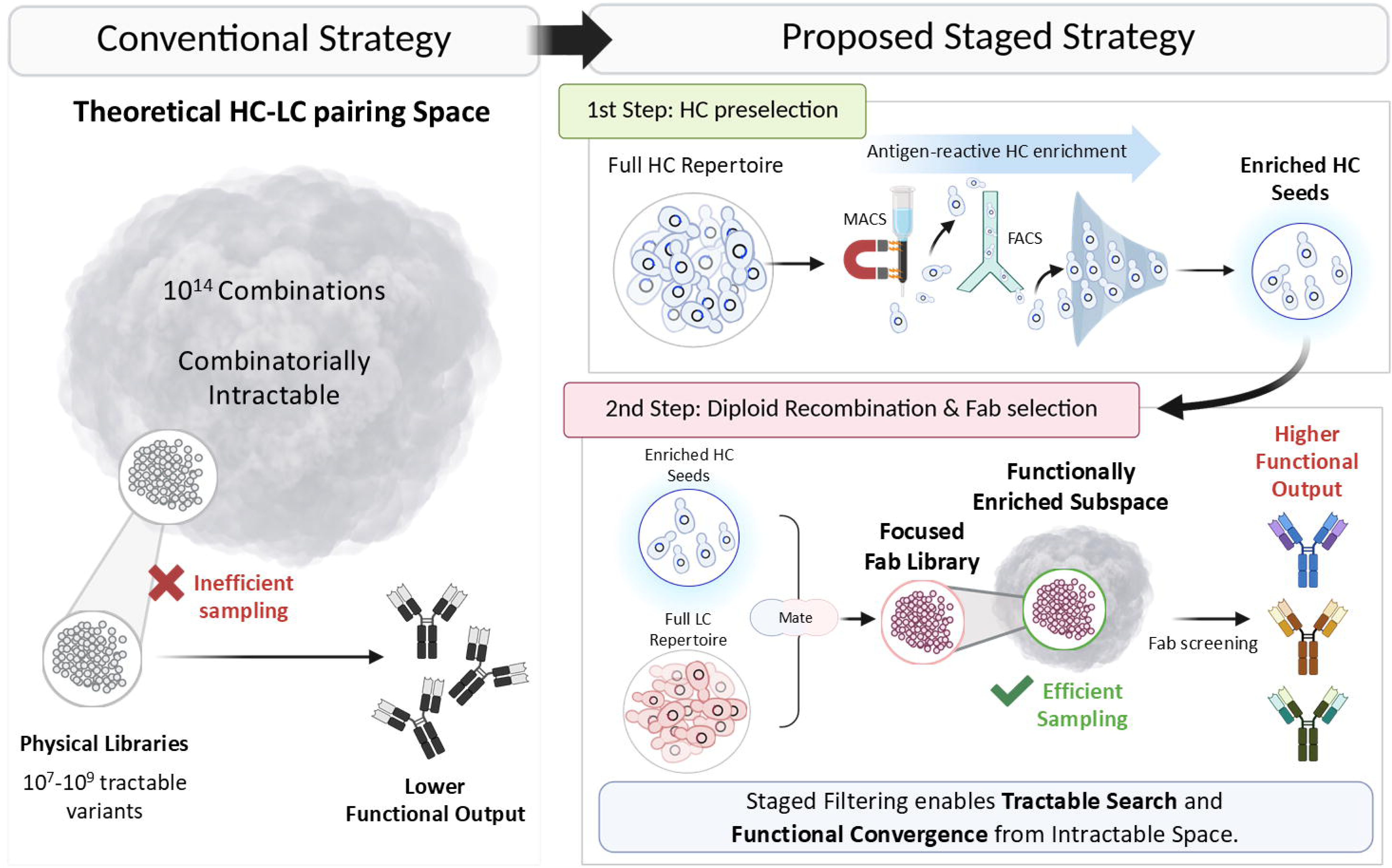

