## Supplementary Data for "Staged heavy-chain filtering enables Fab discovery from combinatorially intractable library spaces"

**Supplementary Figure Legends**

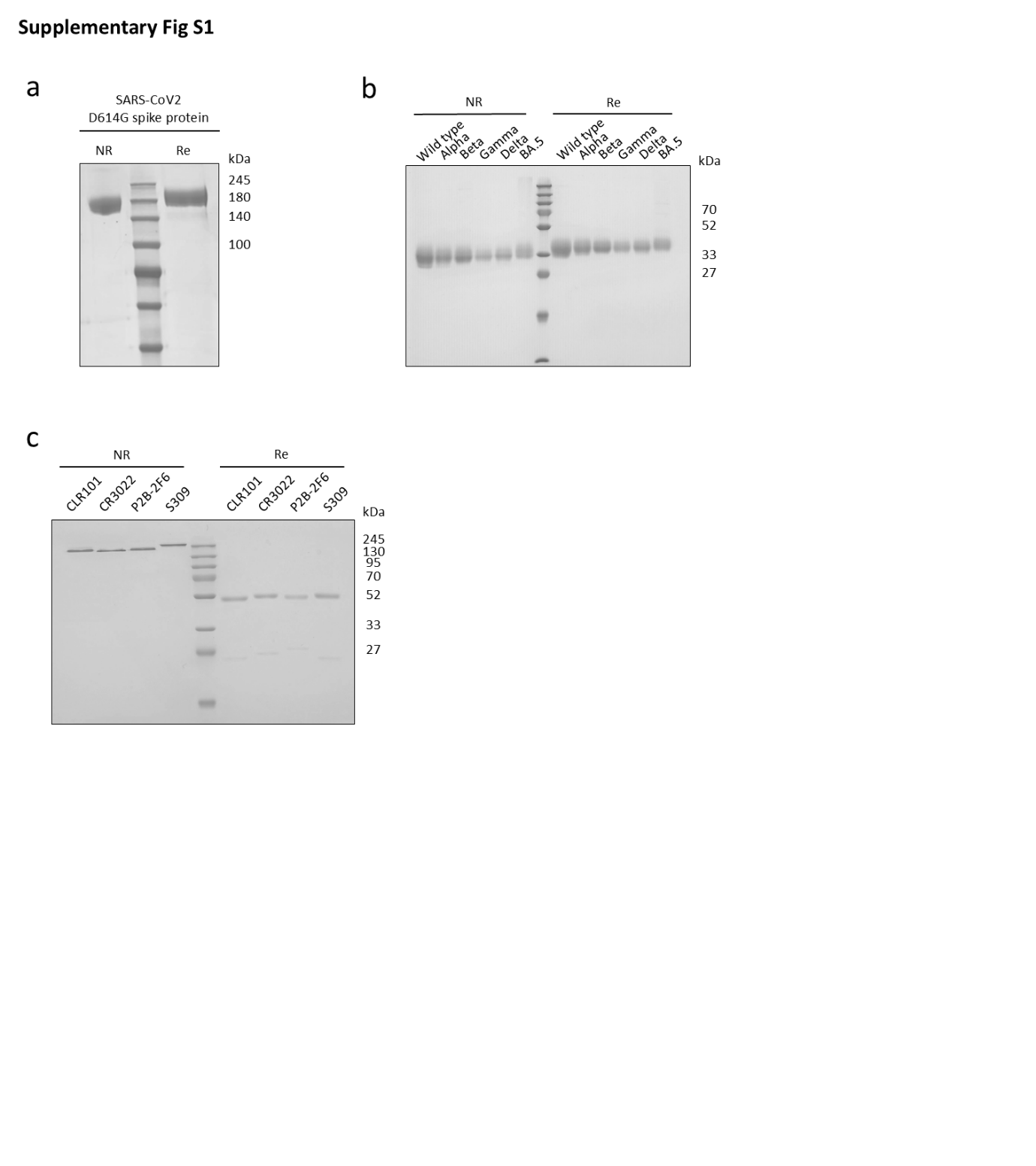

**Supplementary Figure S1. SDS-PAGE analysis of recombinant proteins and antibodies.**

(a) Purified SARS-CoV-2 D614G spike protein. (b) Purified SARS-CoV-2 variant RBD antigens. (c) Purified CLR101 and reference antibodies (CR3022, P2B-2F6, and S309). NR and Re denote non-reducing and reducing conditions, respectively.

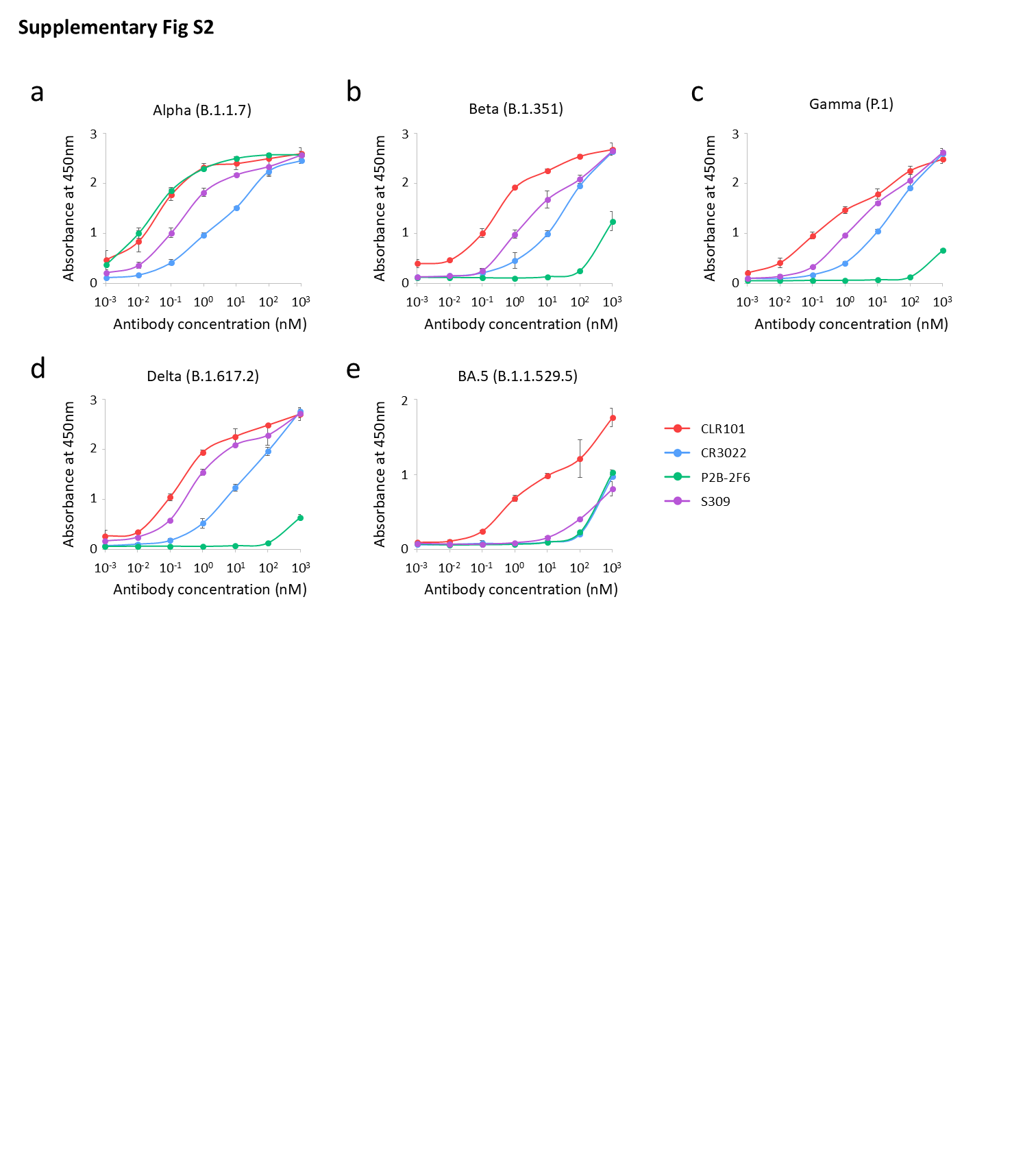

**Supplementary Figure S2. Binding analysis of antibodies against SARS-CoV-2 variant RBDs.** (a–e) Concentration-dependent ELISA binding curves of CLR101, CR3022, P2B-2F6, and S309 against Alpha, Beta, Gamma, Delta, and BA.5 RBDs (n = 2 independent experiments, mean ± s.d.).

| VH PCR | |
| --- | --- |
| Forward (5’ to 3’) | Reverse (5’ to 3’) |
| GATAAAAGAGAGGCCGCTAGGGCCCAGRTGCAGCTGGTGSARTCTGG | TGAGGAGACGGTGACCAGGGTGCCCGATGGGCCCTTGGTGGAGGC |
| GATAAAAGAGAGGCCGCTAGGGCCSAGGTCCAGCTKGTRCAGTCTGG | TGAAGAGACGGTGACCATTGTCCCCGATGGGCCCTTGGTGGAGGC |
| GATAAAAGAGAGGCCGCTAGGGCCCAGRTCACCTTGAAGGAGTCTG | TGAGGAGACGGTGACCAGGGTTCCCGATGGGCCCTTGGTGGAGGC |
| GATAAAAGAGAGGCCGCTAGGGCCGAGGTGCAGCTGKTGGAGWCY | TGAGGAGACGGTGACCGTGGTCCCCGATGGGCCCTTGGTGGAGGC |
| GATAAAAGAGAGGCCGCTAGGGCCSARGTGCAGCTGGTGSAGTCTGG |  |
| GATAAAAGAGAGGCCGCTAGGGCCTCAACACAACGGTTCCCAGTTA |  |
| GATAAAAGAGAGGCCGCTAGGGCCCAGSTGCAGCTGCAGGAGTCSG |  |
| GATAAAAGAGAGGCCGCTAGGGCCCAGGTGCAGCTACAGCAGTGGG |  |
| GATAAAAGAGAGGCCGCTAGGGCCCAGGTACAGCTGCAGCAGTCA |  |

| Vk PCR | |
| --- | --- |
| Forward (5’ to 3’) | Reverse (5’ to 3’) |
| GATAAAAGAGAGGCCGCTAGGGCCGACATCCRGDTGACCCAGTCTCC | TTTGATYTCCASCTTGGTCCCCTGACAGATGGTGCAGCCACCGTACG |
| GATAAAAGAGAGGCCGCTAGGGCCGATATTGTGMTGACBCAGWCTCC | TTTGATATCCACTTTGGTCCCCTGGCACAGATGGTGCAGCCACCGTACG |
| GATAAAAGAGAGGCCGCTAGGGCCGAWRTTGTGMTGACCCACACTCC | TTTAATCTCCAGTCGTGTCCCCTGCACAGATGGTGCAGCCACCGTACG |
| GATAAAAGAGAGGCCGCTAGGGCCGAAATTGTRWTGACRCAGTCTCC |  |
| GATAAAAGAGAGGCCGCTAGGGCCGAAACGACACTCACGCAGTCTC |  |

**Supplementary Table S1.** List of primer sequences for construction of the human BCR library.

|  | HCDR1 | HCDR2 | HCDR3 |
| --- | --- | --- | --- |
| HC1 | SYYWS | EIYHNGSSNYNPSLKS | CGDCYPYYGMDV |
| HC2 | SGAYYWS | YIYHSGSAYYNPSLKS | CSGVNEDAVDV |
| HC3 | GYHWT | EINHSGSTNYSPSLKS | GHTRSVRGVIPFDY |
| HC4 | SISSAWS | RTYYRSKWYYDYAVSVKG | GLPSLTRGGLDP |
| HC5 | SGSYYWG | SIFYTGATYYNPSLKS | GRRTHTVVVTAPLDY |
| HC6 | GYYCI | EINHGGSTNYNPSLKS | GGQDSTLYSCFQH |
| HC7 | SYYWS | EINHSGSTNYNPSLEG | PHLYYDKKGKAFDI |
| HC8 | GYYWS | EITHGGITNYNPSLKS | GGAVYSSAFFDY |
| HC9 | APYWG | EINHSGSTDYNPSLKS | GEGVGWSPNANNNYYYGMAV |
| HC10 | SYYWN | YIYYSGTSYYNPSLKS | PLLESSGPSHYHSGLDI |
| HC11 | KYYWS | EINQGGITNYNPSLKS | MALPPFDY |
| HC12 | DYYWS | EINQIGITRYNPSLPS | GTTDVNMVIVLIGMIYYLDL |
| HC13 | GYYWS | EVDHSGSSTYNPSLKS | GIADMTMVVLVITGKSHAFD |
| HC14 | SYGMY | IIWHDGSNRNYTDSVKG | FYSSRHGMDV |

|  | LCDR1 | LCDR2 | LCDR3 |
| --- | --- | --- | --- |
| LC1 | RASQTISSWLA | DASNRAT | QQSYSRRT |
| LC2 | RASQSVGSDLA | GASTRAT | QQYNDWPPF |
| LC3 | QASQDIRNHLN | KASGLES | QQYNNWPPWT |
| LC4 | MASQNVSRNLA | AKSTRAT | QQYGSSLIT |
| LC5 | RASENINHYLN | AASSLQS | QQTYISPPT |
| LC6 | RASENINHYLN | AASSLQS | QQSYSTPRS |
| LC7 | KSSQSVLYSSNNKNYLA | WASTRES | QQYYNTPPLT |
| LC8 | QASQDISNYLN | AASSLQS | QQYGSSPPT |
| LC9 | RASQGISHYLS | AASSLQS | QQYHTFPYT |
| LC10 | KSSQSLFYSSNNRNYLT | GASTRDS | HQYYSTPYT |
| LC11 | RASQSVSSSYLA | GASSRAT | QQYYSFPYT |

**Supplementary Table S2.** Sequence analysis of the initial heavy-chain (HC) and light-chain (LC) libraries generated from PBMCs of patients with COVID-19, showing the diversity of complementarity-determining region (CDR) sequences.

| VH | VH-FW1 | VH-CDR1 | VH-FW2 | VH-CDR2 | VH-FW3 | VH-CDR3 | VH-FW4 |
| --- | --- | --- | --- | --- | --- | --- | --- |
| CLR101 | QVQLVQSGGGLVQPGRSLRLTCVASGFTFD | DYAMH | WVRQAPGRGLEWVA | VVTWNSGTIGYADSVKG | RFIIIRDNAANSLYLQMNSLTAEDTAIYYCAK | DISGLLRFGGERYAFDV | WGQGTMVTVSS |
| CLR102 | QVQLVQSGGGLVQPGRSLRLTCVASGFTFD | DYAMH | WVRQAPGRGLEWVA | VVTWNSGTIGYADSVKG | RFIIIRDNAANSLYLQMNSLTAEDTAIYYCAK | DISGLLRFGGERYAFDV | WGQGTMVTVSS |
| CLR103 | QVQLVQSGGGLVQPGRSLRLTCVASGFTFD | DYAMH | WVRQAPGRGLEWVA | VVTWNSGTIGYADSVKG | RFIIIRDNAANSLYLQMNSLTAEDTAIYYCAK | DISGLLRFGGERYAFDV | WGQGTMVTVSS |

| VL | VL-FW1 | VL-CDR1 | VL-FW2 | VL-CDR2 | VL-FW3 | VL-CDR3 | VL-FW4 |
| --- | --- | --- | --- | --- | --- | --- | --- |
| CLR101 | DIQVTQSPSSLSASVGDRVTITC | RASQSVSSYLA | WYQQKPGKAPKLLIY | GASSRAT | GIPDRFSGSGSGTDFTLTISRLEPEDFAIYYC | QQRET | FGQGTKVEIK |
| CLR102 | DIQVTQSPSSLSASVGDRVTITC | RASQSISNYLN | WYQQKPGKAPKLLIY | SASSLQR | GVPSRFSGSGSGTDFTLTISSLHPEDFATYYC | QQSYGPPVT | FGQGTKVDIK |
| CLR103 | DIQVTQSPSSLSASVGDRVTITC | RASQSISSYLN | WYQQKPGKAPKLLIY | SASTLQS | GVPSRFSGSGSGTDFTLTIDSLQAEDFATYYC | QQAYSVPIT | FGQGTKVDIK |

**Supplementary Table S3.** Complete amino acid sequences of representative SARS-CoV-2-targeting Fab clones CLR101, CLR102, and CLR103.

|  | EC_50_ (nM) | | | | |
| --- | --- | --- | --- | --- | --- |
|  | RBD alpha | RBD beta | RBD gamma | RBD delta | RBD BA.5 |
| CLR101 | 0.047 ± 0.004 | 0.367 ± 0.033 | 0.459 ± 0.124 | 0.215 ± 0.005 | 26.9 ± 0.5 |
| CR3022 | 5.12 ± 0.28 | 38.6 ± 9.5 | 33.9 ± 5.7 | 40.2 ± 21.5 | >1000 |
| P2B-2F6 | 0.025 ± 0.001 | >1000 | >1000 | 227.7 ± 47.3 | >1000 |
| S309 | 0.193 ± 0.006 | 7.72 ± 1.87 | 6.11 ± 0.14 | 0.853 ± 0.017 | 302.0 ± 18.3 |

**Supplementary Table S4.** Apparent half-maximal effective concentration (EC₅₀) values of CLR101 and reference anti-RBD antibodies against SARS-CoV-2 variant RBDs (n = 2 independent experiments, mean ± s.d.).

| T Onset (°C) | Error | Tm1 (°C) | Error | Tm2 (°C) | Error |
| --- | --- | --- | --- | --- | --- |
| 63.74 | 0.08 | 73.53 | 0.02 | 77.89 | 0.07 |

**Supplementary Table S5.** Thermal stability parameters of CLR101 IgG1 determined by intrinsic differential scanning fluorimetry, also referred to as SUPR-DSF (n = 3 independent replicates).
